# Beta-Hydroxybutyrate Attenuates Bronchial Smooth Muscle Pro-Inflammatory Cytokine Production

**DOI:** 10.1101/2025.02.19.639048

**Authors:** V. Amanda Fastiggi, Madeleine M. Mank, Matthew E. Poynter

## Abstract

Asthma is a common airway condition causing breathing difficulties due to reversible airflow obstruction. It often affects obese individuals, with symptoms triggered by environmental factors that induce immune responses, leading to inflammation and bronchoconstriction. Bronchial smooth muscle (BSM) plays a central role in airway narrowing, driven by type 2 immune responses involving cytokines like IL-4, IL-5, and IL-13, along with leukocytes including eosinophils and type 2 T-helper cells. These responses cause structural changes such as fibrosis and airway thickening, while BSM cells worsen asthma by releasing pro-inflammatory cytokines in response to allergens, microbial signals, or inflammatory cytokines from other cells. While current treatments manage asthma in most patients, alternative therapies are needed for difficult-to-treat cases, particularly prevalent in obese, allergic individuals. Emerging research suggests that therapeutic ketosis, induced by dietary changes or ketone supplementation, may reduce airway hyperresponsiveness and inflammation. The primary ketone body, β-hydroxybutyrate (BHB), produced during carbohydrate scarcity, acts via cell-surface receptors and transporters, potentially mitigating asthma symptoms. Weight loss and caloric restriction increase ketone levels, correlating with reduced inflammation and improved asthma outcomes. We hypothesized that β-hydroxybutyrate (BHB) reduces bronchoconstriction and inflammation in asthma by targeting bronchial smooth muscle. Using human bronchial smooth muscle cells (HBSMC) *in vitro,* we demonstrate herein that BHB suppresses IL-1β-induced pro-inflammatory cytokine production through Free Fatty Acid Receptor 3 (FFAR3) activation. These findings suggest that bronchial smooth muscle is a key target of therapeutic ketosis, supporting BHB’s potential benefits in preclinical asthma models.

## Introduction

Asthma, a condition affecting the airways, is typically provoked by environmental triggers that initiate immune reactions in the lungs, leading to excessive inflammation and bronchoconstriction, and affects approximately 8.9% of adults and 6.7% of children in the United States^1^. Asthma-associated airway hyperresponsiveness can be evaluated using the methacholine challenge test, a diagnostic procedure designed to assess the responsiveness of airway smooth muscle. This test involves the inhalation of methacholine, a substance that causes airway constriction in individuals with asthma or other airway hyperreactivity conditions. By monitoring the degree of airway narrowing in response to methacholine, the test helps healthcare providers evaluate the functionality of the airway smooth muscle and plays a crucial role in confirming the diagnosis of asthma^2,3^. Bronchial smooth muscle is central to the pathophysiology of asthma, as its contraction causes airway narrowing, resulting in obstruction and breathing difficulties^4^.

Bronchial smooth muscle has been implicated in the pathophysiology of bronchial inflammation both as a target as well as a mediator of inflammatory reactions^5,6^. Type 2 inflammation is most common in asthmatics and is associated with the cytokines interleukin(IL)-4, IL-5, and IL-13, inflammatory cells including eosinophils, mast cells, basophils, type 2 innate lymphoid (ILC2) and T helper (Th2) lymphocytes, and immunoglobulin E (IgE)-producing plasma cells^7,8^. These asthma-relevant cell types can release pro-inflammatory mediators such as cytokines and chemokines that can affect bronchial smooth muscle cells^6,9,10^, the products of which further exacerbate airway hyperresponsiveness and remodeling^4,5,10–12^. Over time, this chronic inflammation can lead to structural changes in the airways, including thickening of the airway walls, fibrosis, and increased mucus production, which in aggregate contribute to the persistent symptoms of asthma^11,13^. We and others have previously reported that bronchial smooth muscle cells secrete pro-inflammatory cytokines in response to immune stimulants, including lipopolysaccharide (LPS) and house dust mite (HDM) extract, as well as cytokines such as IL-1β^10,14^ produced from the other asthma-relevant cell types. IL-1β is a potent pro-inflammatory cytokine produced from both immune (leukocytes) and non-immune cells^12,15^. Importantly, IL-1β is elevated in the airways^16^ and serum of allergic and non-allergic asthmatic subjects compared to controls^17^.

Current treatments for asthma include broncho-relaxing β-agonists, anti-inflammatory corticosteroids, and biological immunotherapies targeting the type 2 immune response^18^. While these therapies provide effective disease control for most patients, individuals with ’difficult-to-treat’ asthma often require alternative or additional approaches, especially as severe asthmatics with type 2 inflammation exhibit poor response to corticosteroids^19^. Recently, we have demonstrated the beneficial effects of increasing circulating ketone bodies, known as therapeutic ketosis, in multiple mouse models of obesity-associated^20^ and allergic asthma^21^.

Ketone bodies, β-hydroxybutyrate (BHB) and acetoacetate (AcAc), are synthesized in the liver from fatty acids^22,23^, either through dietary modifications^23,24^ or the mobilization of adipose tissue during periods of increased energy demand^23^. Once produced, these ketone bodies are transported via the bloodstream to cells throughout the body.

Ketone bodies have been implicated in modulating key pathological processes in asthma. Initially recognized as an energy substrate for ATP production in the Krebs cycle during carbohydrate scarcity, ketone bodies have since been shown to influence cellular processes through various mechanisms. They can stimulate cell-surface receptors such as the G-protein coupled receptors Hydroxycarboxylic Acid Receptor 2 (HCAR2 or GPR109a) and Free Fatty Acid Receptor 3 (FFAR3 or GPR41)^23,25–29^, or by uptake through transporters such as Monocarboxylate Transporter 1 (MCT1)^27,30^. Beyond these roles, ketone bodies also function as antioxidants^31,32^ and exert effects through transcriptional and epigenetic regulation^22,27,33^, including suppression of nuclear factor-κB (NF-κB) activation^31,25^. Furthermore, they possess anti-inflammatory properties, notably through inhibition of the NLRP3 inflammasome, which decreases IL-1β production^34–36^. *In vivo*, dietary interventions such as weight loss and alternate-day caloric restriction raise BHB levels, which correlate with reduced asthmatic symptoms, including lower oxidative stress and inflammation in obese asthmatic patients^14,37,38^. Early ketone body elevations are linked to improved asthma symptoms in obese patients following bariatric surgery^39,40^, alternate-day caloric restriction^37^, and during treatment with GLP-1R agonists^41^. Independent of weight loss, therapeutic ketosis can be achieved through providing exogenous ketones or ketogenic precursors (*e.g.,* ketone esters). Therapeutic ketosis can be induced by providing exogenous ketones or ketogenic precursors (e.g., ketone esters). Notably, increasing ketone levels is generally well-tolerated in humans^42,43^, highlighting its potential as a viable therapeutic approach.

Therapeutic ketosis achieved through feeding a high fat/low carbohydrate diet or by supplementing the normal diet with ketone esters, significantly and substantially decreases methacholine hyperresponsiveness in mouse models of asthma^20,21^ and also decreases methacholine responsiveness in non-asthmatic mice^20^. As methacholine functions through the activation of bronchial smooth muscle^44–47^, we sought to study these cells more directly to explore the impact of ketone bodies on the ability of bronchial smooth muscle to contribute to the inflammatory environment in asthma. We have established a well-functioning *in vitro* model using human bronchial smooth muscle cells (HBSMC), allowing us to directly observe the effects of BHB on this cell type^21^. Herein, we used HBSMC as an *in vitro* model of bronchial smooth muscle to address gaps and provide a valuable platform for investigating the therapeutic potential of ketone bodies and their underlying mechanisms of action.

We hypothesized that BHB can mitigate asthma-related pathologies, in part, by inhibiting pro-inflammatory cytokine production from bronchial smooth muscle cells. Our objectives were to assess the effectiveness of BHB in reducing IL-1β-induced pro-inflammatory cytokine secretion *in vitro* and to identify the mechanisms by which BHB may influence these effects.

Gaining a deeper understanding of the efficacy and mechanisms by which BHB reduces pro-inflammatory cytokine production by human bronchial smooth muscle *in vitro* could offer valuable insights into potential novel targets for treating asthma and its associated symptoms.

## Materials and Methods

### Study Approval

Studies involving potentially hazardous material were reviewed and approved by the University of Vermont’s Institutional Biosafety Committee (REG201900052).

### Human Bronchial Smooth Muscle Cell Culture

Primary human bronchial smooth muscle cells (HBSMC) isolated from a 45-yr-old female patient with asthma (Lonza, Morristown, NJ, Lot No. 00194850, Batch No. 0000195154) were cultured in smooth muscle cell growth medium-02 BulletKit (Lonza, CC-3182) according to the manufacturer’s instructions at 37°C in 95% humidified air containing 5% CO_2_. The cells were used within the first seven passages to ensure proper smooth muscle phenotype. Cell authentication was performed by Lonza (negative Factor VIII-related antigen, positive α-Actin expression) and cells tested negative for mycoplasma (MycoDect Mycoplasma Detection Kit (Alstem, Richmond, CA)) before being utilized for experiments. For agonist-induced cytokine production experiments, human bronchial smooth muscle cells (HBSMC) were stimulated with 50 ug/mL House Dust Mite extract (HDM) (Stagrallery/Greer, Cat No.B70, Lot No.390992), 100 ng/mL ultrapure lipopolysaccharide (LPS) from Escherichia coli 0111:B4 (Invivogen, Cat No.tlrl-3pelps), or 10 ng/mL recombinant human IL-1β (Stem Cell Technologies, Cat No.78034.1).

For simultaneous exposure and stimulation experiments, HBSMC were plated at 5×10^4^ cells/cm^2^ in 1mL of media in a 12-well plate and allowed to grow for 48 hours at 37°C and 5% CO_2_.

HBSMC were exposed for 24 hours with vehicle or 1.25-10 mM beta-hydroxybutyric acid (BHBA) (Sigma-Aldrich, St. Louis, MO, Cat No.166898), sodium beta-hydroxybutyrate (NaBHB)(Sigma-Aldrich, Cat No.54965), (R)-beta-hydroxybutyric acid ((R)-BHBA) (Sigma-Aldrich, Cat No.54920), (S)-beta-hydroxybutyric acid ((S)-BHBA) (Sigma-Aldrich, Cat No.54925), FFAR3 agonist, AR420626 (AR) (Caymen, Cat No.17531), or MCT1 inhibitor, AZD3965 (Med Chem Express, Cat No.HY-12750), butyric acid (BA), sodium butyrate (SB), nicotinic acid (NA), and sodium nicotinic acid (NaNA) while being stimulated with 10 ng/mL recombinant human IL-1β (Stem Cell Technologies, Cat No.78034.1).

In pre-exposure experiments with subsequent stimulation, HBSMC were plated at 5×10^4^ cells/cm^2^ in 1mL of media in a 12-well plate and allowed to grow for 24 hours at 37°C and 5% CO_2_. After the initial 24 hours, cells were exposed to vehicle or 1.25-10 mM BHBA, NaBHB, (R)- BHBA, (S)-BHBA, or AR for another 24 hours before being washed and stimulated with 10 ng/mL recombinant human IL-1β.

### Cytokine Immunoassays

Conditioned medium from the cell culture studies was collected at the indicated time points, centrifuged for 10 minutes at 3300 x *g* to eliminate debris, transferred into new tubes or multi-well plates, and frozen at -20°C until analysis. ELISAs to quantitate human IL-8/CXCL8 and IL-6 levels (DuoSets from R&D Systems) were used according to the manufacturer recommendations, with samples diluted to coincide with the range of the standards. A custom human magnetic Luminex assay (R&D Systems, Minneapolis, MN) was performed per manufacturers’ instructions to measure cytokines present in the HBSMC supernatants, including CCL2, CCL4, CCL20, CXCL1, CXCL2, and G-CSF. Luminex assays were performed using the Bio-Rad Bio-Plex suspension array system, Bio-Rad Bio-Plex Pro II wash station, and Bio-Plex Manager 6.0 software (Bio-Rad, Hercules, CA). Concentrations were calculated by five-place logistic regression from standards within 70%-130% of the expected values using Bio-Rad Manager 6.0. Studies were conducted with all BHB compounds used in the experiments to validate that they did not interfere substantially with the ability of the ELISAs to accurately quantify the recombinant standards.

### Statistical Analyses

All experiments included multiple biological replicates for each condition. Outliers were identified and removed using the ROUT method (Q=1%) and the cleaned data were analyzed using unpaired one-way ANOVA with post-hoc multiple comparisons: Tukey’s test (for comparing all means) or Dunnett’s test (for comparing each mean to a control), were performed using GraphPad Prism 10.2.3 (GraphPad Software, Inc., La Jolla, CA). Data are presented as means ± SEM from representative experiments. P values below 0.05 are considered statistically significant, and significance levels are indicated in the figure legends.

## Results

### Human Bronchial Smooth Muscle Cells (HBSMC) Inducibly Secrete Pro-Inflammatory Cytokines

Although known primarily as a contractile cell type, bronchial smooth muscle can produce many chemokines and cytokines in response to various agonists^10,14,48^. A custom human magnetic Luminex assay was performed to measure cytokines present in the HBSMC supernatants following stimulation for 24 hours with House Dust Mite extract (HDM), lipopolysaccharide (LPS), and human recombinant IL-1β to determine which pro-inflammatory cytokines were produced. This analysis revealed that CCL2 **(Figure 1A)**, CCL4 **(Figure 1B)**, CCL20 **(Figure 1C)**, CXCL1 **(Figure 1D)**, CXCL2 **(Figure 1E)**, G-CSF **(Figure 1F)**, IL-6 **(Figure 1G)**, and CXCL8/IL-8 **(Figure 1H)** were produced significantly when HBSMC were stimulated with 10 ng/mL of recombinant human IL-1β.

**Figure 1.**
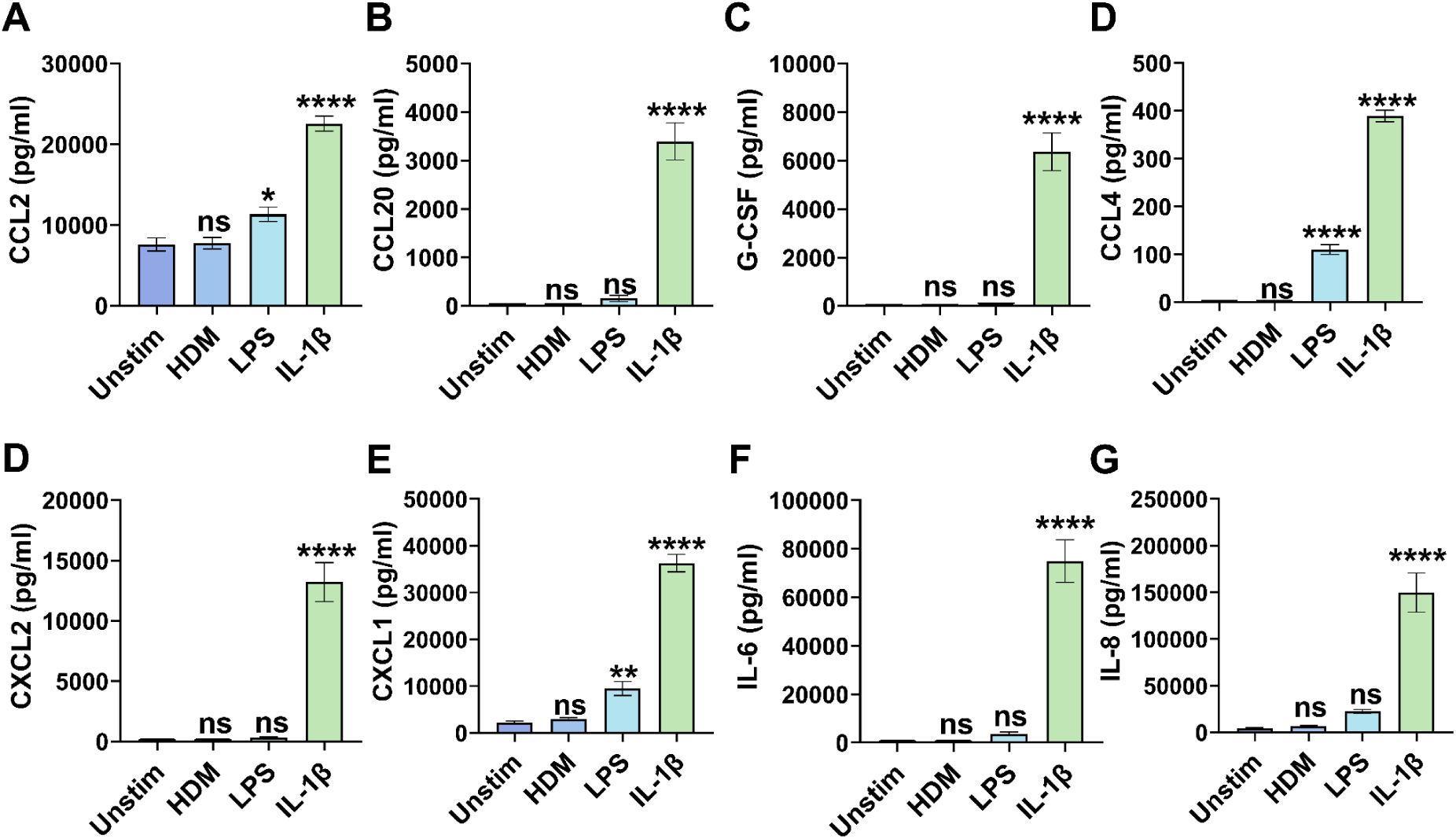
IL-1β induces pro-inflammatory cytokine secretion from HBSMC. HBSMCs were unstimulated (Unstim) or stimulated *in vitro* with 50 mg/mL House Dust Mite extract (HDM), 100 ng/mL lipopolysaccharide (LPS), or 10 ng/mL recombinant human IL-1β for 24 hours and **(A)** CCL2, **(B)** CCL20, **(C)** G-CSF, **(D)** CCL4, **(E)** CXCL2, **(F)** CXCL1 **(G)** IL-6 and **(H)** CXCL8/IL-8 levels were measured. ∗*P* ≤ 0.05, ∗∗*P* ≤ 0.01, ∗∗∗*P* ≤ 0.001, ∗∗∗∗*P* ≤ 0.0001 compared to the vehicle.

### BHB Attenuates IL-1β-Induced Pro-Inflammatory Cytokine Production From HBSMC

Ketone bodies, acetoacetate (AcAc) and especially β-hydroxybutyrate (BHB), have been reported to exert anti-inflammatory effects, especially via inhibition of NLRP3 inflammasome activation^34–36^. In our recent studies, we demonstrated the therapeutic benefits of elevating circulating ketone bodies—a state known as therapeutic ketosis—in multiple mouse models of both obese^20^ and allergic asthma^21^, which effectively mitigated methacholine-induced airway hyperresponsiveness and significantly improved lung function *in vivo*^20,21^. We also demonstrated *in vitro* that BHB attenuated pro-inflammatory cytokine secretion from several asthma-relevant cell types, including macrophages, CD4 T lymphocytes, and airway epithelium^21^. Building on these findings, we investigated the effects of ketone bodies on IL-1β-induced pro-inflammatory cytokine production from human bronchial smooth muscle cells (HBSMC). We determined that simultaneous exposure for 24 hours to beta-hydroxybutyric acid (BHBA) or sodium beta-hydroxybutyrate (NaBHB) in the presence of 10ng/mL human recombinant IL-1β caused a dose-dependent decrease in the concentrations of the pro-inflammatory cytokines IL-6 **(Figure 2A)** and CXCL8/IL-8 **(Figure 2B)** compared to cells stimulated with IL-1β in the absence of BHB or in the presence of appropriate pH controls. BHBA significantly decreased both IL-6 and IL-8 concentrations in a dose-dependent manner, but when neutralized to a pH of 7 the robust effect in attenuating IL-6 secretion at all concentrations and in attenuating IL-8 secretion at a BHBA concentration of 10mM was lost. NaBHB did not exert a robust inhibitory effect on pro-inflammatory cytokine production until the pH of the 10mM working solution was matched to the pH of the working concentration of 10mM BHBA. The acidified NaBHB exerted a more robust inhibitory dose response than the non-acidified NaBHB or the neutralized BHBA. An equivalent concentration of HCl matching the acidity (pKa) of the acidic BHB molecules (10mM) modestly attenuated IL-1β-induced secretion of IL-6 and IL-8, but not to the extent of the BHBA or the acidified NaBHB.

**Figure 2.**
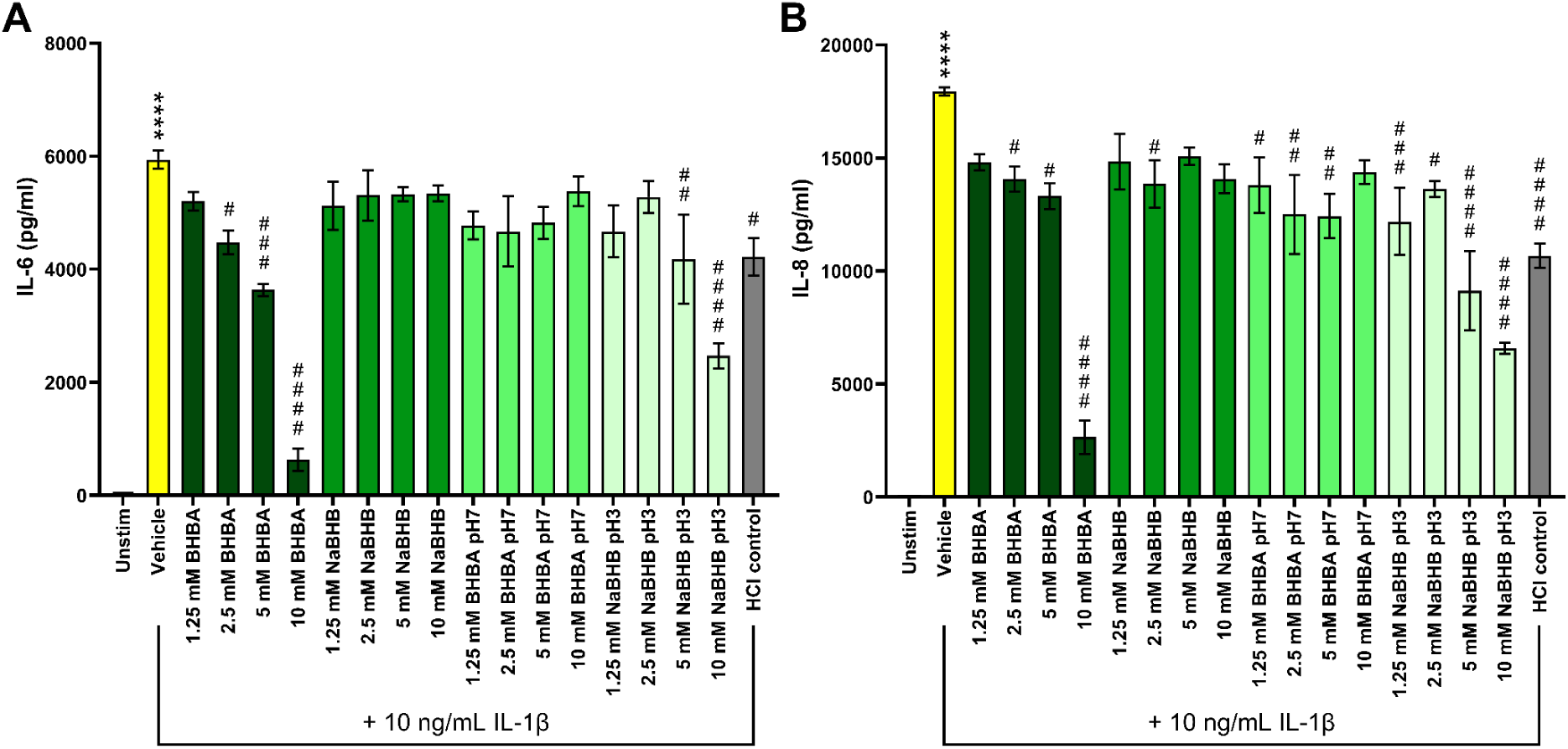
BHB dose-dependently attenuates IL-1β-stimulated pro-inflammatory cytokine production from HBMSC. HBSMCs were unstimulated (Unstim) or stimulated *in vitro* with 10 ng/mL recombinant human IL-1β in the presence of increasing concentrations of BHBA, NaBHB, or the appropriate pH controls of each for 24 hours and **(A)** IL-6 or **(B)** CXCL8/IL-8 were measured. n=4 per group; values are inclusive of studies performed three times. ∗∗∗∗*P* ≤ 0.0001 compared to the vehicle, #*P* ≤ 0.05, ##*P* ≤ 0.0, ###*P* ≤ 0.001, ####*P* ≤ 0.0001 compared to IL-1β.

### Inhibition of Pro-Inflammatory Cytokine Production from HBSMC by BHB Is Not Enantioselective

To determine if the inhibitory effect elicited by BHB on IL-1β-induced IL-6 and IL-8 production was through a mechanism dependent on the metabolism of BHB as an energetic substrate, we utilized (R)-beta-hydroxybutyric acid ((R)-BHBA) and (S)-beta-hydroxybutyric acid ((S)-BHBA), the individual enantiomers of the racemic BHBA, as well as included exposure to the (R) enantiomer of NaBHB, and a cocktail representative of circulating ketone bodies *in vivo*. The BHB cocktail consists of 78% BHB, 20% AcAc, and 2% acetone, which are biologically consistent with the concentrations of ketone bodies in circulation during fasting^49^. In this study, with simultaneous exposure to the BHB compounds and 10 ng/mL IL-1β for 24 hours, we determined that both the (R) and (S) enantiomers, the (R) enantiomer of NaBHB, and the BHB cocktail significantly inhibited IL-1β-induced secretion of IL-6 **(Figure 3A)** and CXCL8/IL-8 **(Figure 3B)** from HBSMC. In this study, all conditions except the (R) enantiomer of NaBHB demonstrated dose-dependent inhibition of pro-inflammatory cytokine production.

**Figure 3.**
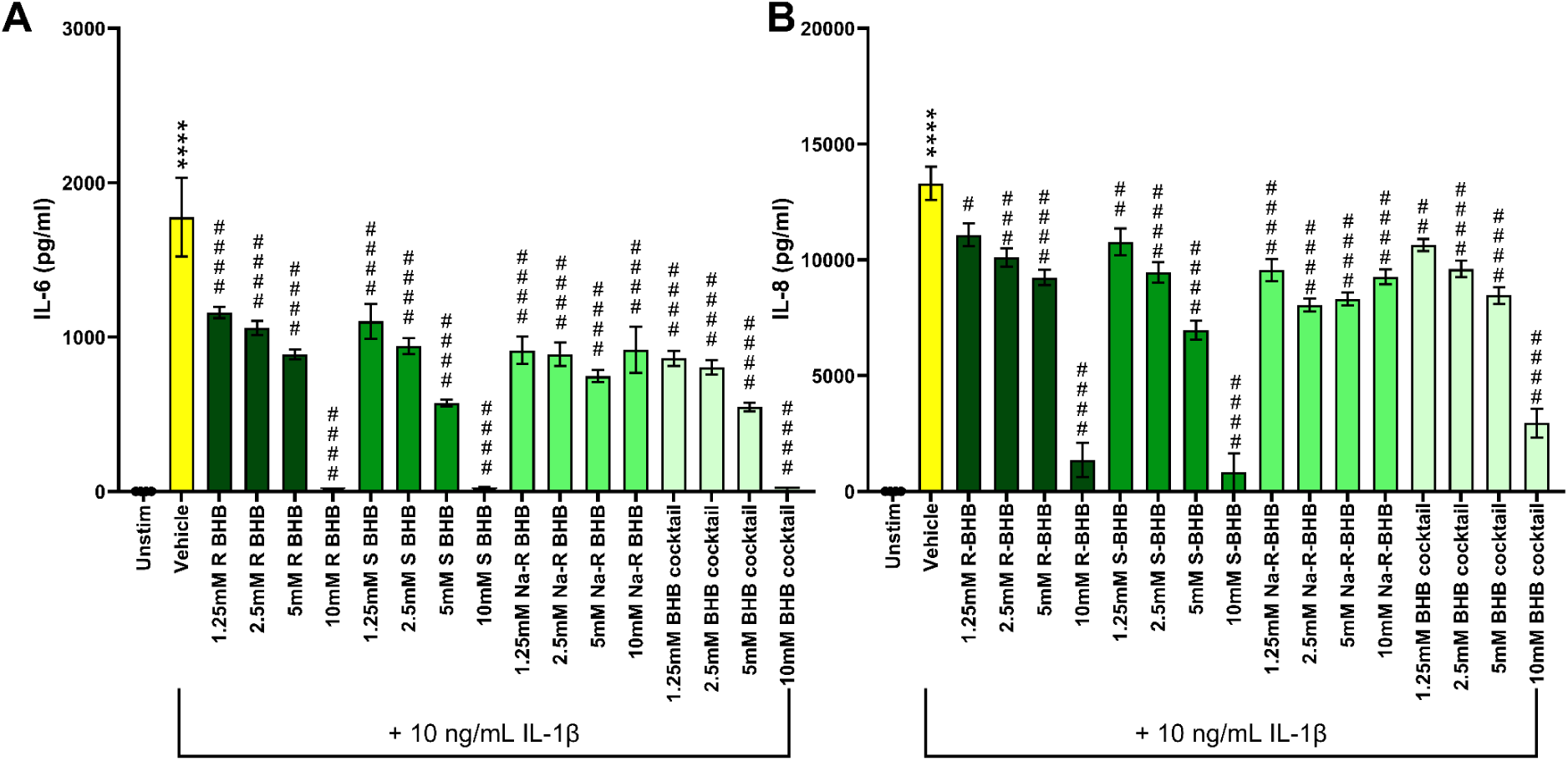
BHB enantiomers dose dependently attenuate IL-1β-stimulated pro-inflammatory cytokine production from HBSMC. HBSMCs were unstimulated (Unstim) or stimulated *in vitro* with 10 ng/mL recombinant human IL-1β in the presence of increasing concentrations of (R)-BHBA, (S)-BHBA, Na-(R)-BHBA, or a BHB cocktail composed of 78% (R)-BHBA, 20% AcAc, and 2% acetone, for 24 hours and **(A)** IL-6 or **(B)** CXCL8/IL-8 were measured. n=4 per group; values are inclusive of studies performed three times. ∗∗∗∗*P* ≤ 0.0001 compared to the vehicle, #*P* ≤ 0.05, ##*P* ≤ 0.01, ###*P* ≤ 0.001, ####*P* ≤ 0.0001 compared to IL-1β.

### Short-Chain Carboxylic Acids Inhibit IL-1β-Induced Pro-Inflammatory Cytokine Production From HBSMC

As BHB has been proposed to elicit effects through activation of cell surface receptors such as hydroxycarboxylic acid receptor 2 (HCAR2)^22^ ^23^ ^25^ ^26^ ^27^, we sought to determine if a pharmacological HCAR2 agonist would elicit similar beneficial inhibitory effects as BHB. Furthermore, as we have previously shown that small molecules structurally similar to BHB have anti-inflammatory effects^34^, and that short-chain carboxylic acids exert anti-inflammatory functions^50,51^, we wanted to determine if butyric acid and sodium butyrate could elicit the same inhibitory effects as BHB. When HBSMC were simultaneously exposed to butyric acid, sodium butyrate, nicotinic acid, and sodium nicotinic acid, as well as the HCAR2 agonist, and stimulated with 10ng/mL IL-1β, production of IL-6 **(Figure 4A)** and CXCL8/IL-8 **(Figure 4B)** was dose-dependently attenuated. Whereas the nicotinic acid and the sodium nicotinic acid both significantly inhibited secretion of the pro-inflammatory cytokines, the butyric acid and sodium butyrate had a much more substantial effect, with the lowest doses of butyric acid and sodium butyrate having a greater impact than the highest dose of exposure, which was 10mM of the nicotinic acid or sodium nicotinic acid.

**Figure 4.**
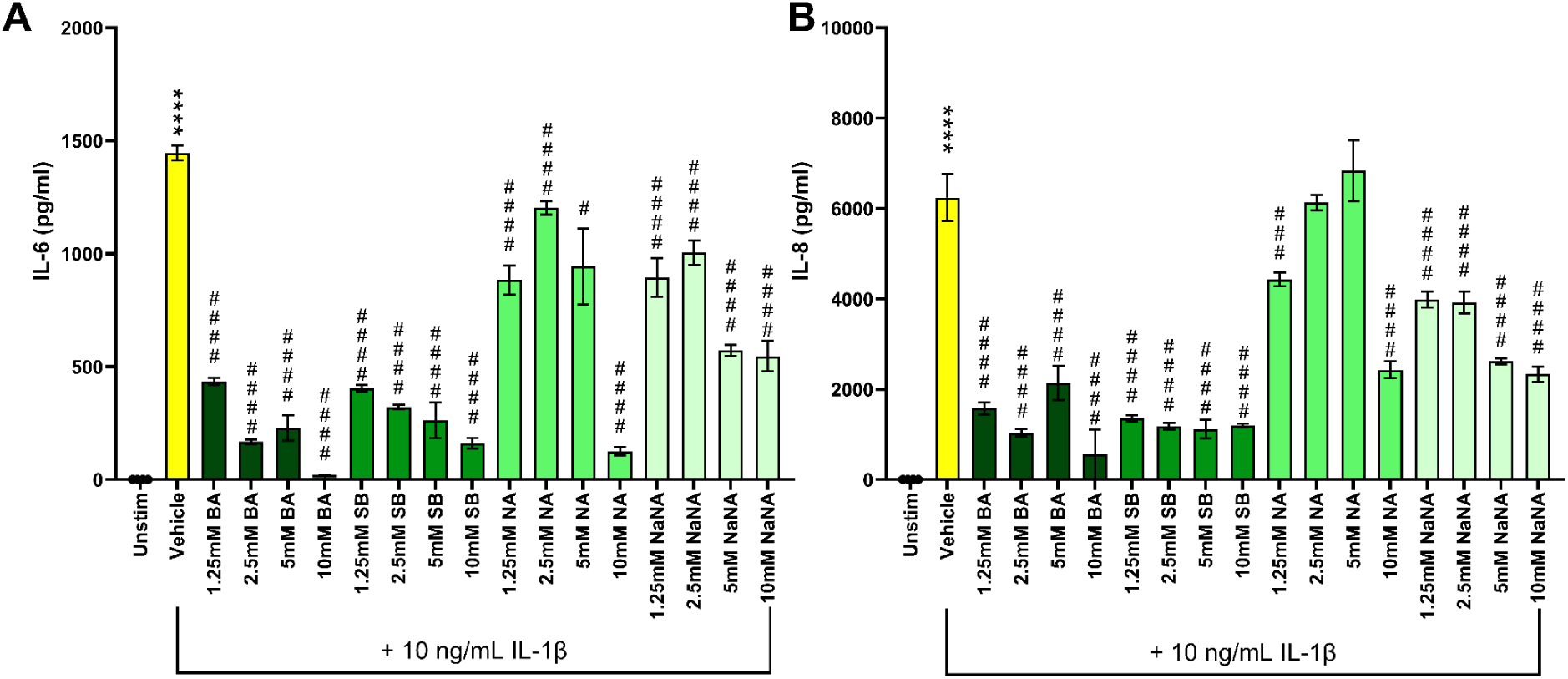
Short-chain carboxylic acids inhibit IL-1β-induced pro-inflammatory cytokine secretion. HBSMCs were unstimulated (Unstim) or stimulated *in vitro* with 10 ng/mL recombinant human IL-1β in the presence of increasing concentrations of butyric acid (BA), sodium butyrate (SB), nicotinic acid (NA), or sodium nicotinic acid (NaNA) for 24 hours and **(A)** IL-6 or **(B)** CXCL8/IL-8 were measured. n=4 per group; values are inclusive of studies performed three times. ∗∗∗∗*P* ≤ 0.0001 compared to the vehicle, #*P* ≤ 0.05, ##*P* ≤ 0.01, ###*P* ≤ 0.001, ####*P* ≤ 0.0001 compared to IL-1β.

### Pre-Exposure of HBSMC to BHBA Attenuates IL-1β-Induced Pro-Inflammatory Cytokine Production

As our previously reported *in vivo* studies have modeled endogenous ketone augmentation through dietary interventions that provide elevated systemic concentrations of these molecules over a protracted period, we sought to model this extended exposure in an *in vitro* system. HBSMC were pre-exposed to biologically relevant concentrations of beta-hydroxybutyric acid (BHBA) for 24 hours, followed by washing and subsequent stimulation with 10 ng/mL human recombinant IL-1β. Pre-exposure to BHBA significantly attenuated IL1β-induced IL-6 **(Figure 5A)** and CXCL8/IL-8 **(Figure 5B)** production in a dose-dependent manner, consistent with the inhibitory effects observed during simultaneous exposure of BHBA and IL-1β. While not as robust as simultaneous stimulation, pretreatment with the higher concentrations of 5mM and 10mM BHBA significantly attenuated IL-6 and IL-8 production from these cells. Although not statistically significant, a trend towards a dose-responsive effect was observed.

**Figure 5.**
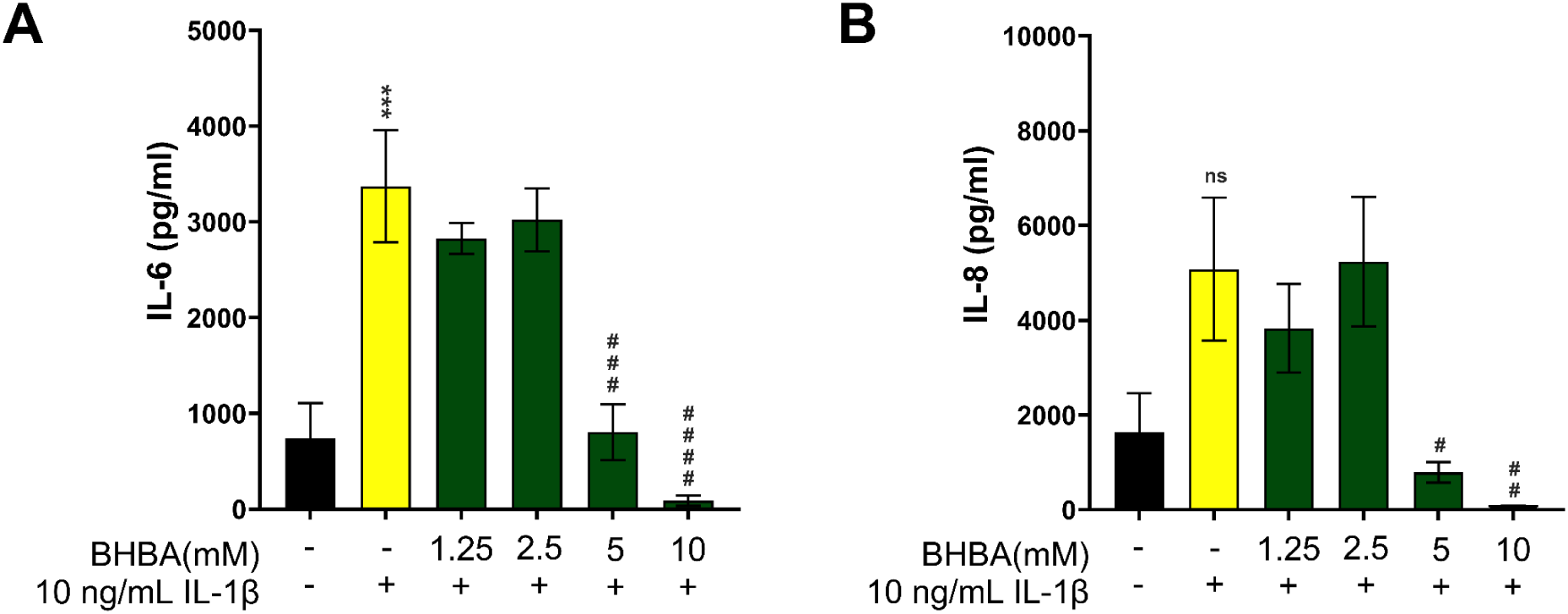
Pre-exposure to BHBA dose-dependently decreases IL-1β-induced pro-inflammatory cytokine secretion. HBSMCs remained unexposed or were exposed *in vitro* to increasing concentrations of beta hydroxybutyric acid (BHBA) for 24 hours, washed and then stimulated with 10 ng/mL recombinant human IL-1β for another 24 hours. **(A)** IL-6 or **(B)** CXCL8/IL-8 were then measured. n=4 per group; values are inclusive of studies performed three times. ∗∗∗*P* ≤ 0.001 compared to the vehicle, #*P* ≤ 0.05, ##*P* ≤ 0.01, ###*P* ≤ 0.001, ####*P* ≤ 0.0001 compared to IL-1β.

### Simultaneous Exposure of BHBA with FFAR3 Agonist Dose-Dependently Attenuates IL-1β-Induced Pro-Inflammatory Cytokine Production

Ketone bodies, particularly BHB, have been proposed to mediate their beneficial effects via uptake through Monocarboxylic Transporter 1 (MCT1)^26,43^ or through a Free Fatty Acid 3 (FFAR3)-dependent pathway^26–28,52^. To evaluate the involvement of these receptors, we tested the MCT1 inhibitor AZD3965 (AZD) in the presence of the BHB compounds. HBSMCs were exposed to biologically relevant concentrations of BHB to model endogenous ketone augmentation, a combination of both the increasing concentrations of BHB and 50µM of the FFAR3 agonist, or a combination of increasing concentrations of BHB and 40nM MCT1 inhibitor (AZD) while stimulated with 10 ng/mL recombinant human IL-1β for 24 hours. The FFAR3 agonist in combination with BHB dose-dependently attenuated IL-6 **(Figure 6A)** and CXCL8/IL-8 **(Figure 6B)** production, and to a greater extent than BHBA alone. In contrast, the presence of AZD did not block the inhibitory effects of BHB on IL-1β-induced pro-inflammatory cytokine production, indicating that these effects are independent of MCT1-mediated uptake of BHB.

**Figure 6.**
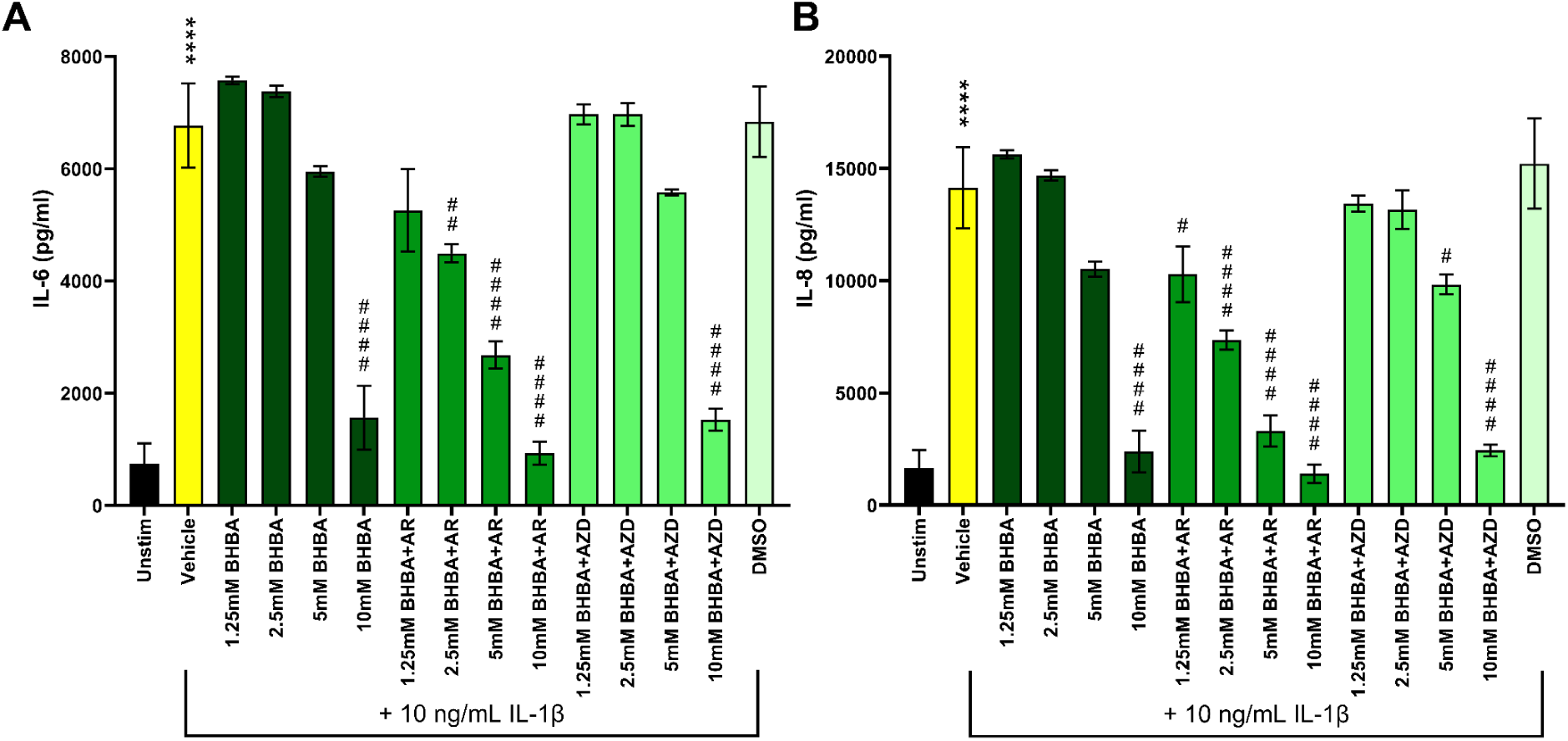
Simultaneous exposure to BHBA with a FFAR3 agonist or MCT1 inhibitor dose-dependently attenuates IL-1β induced pro-inflammatory cytokine secretion. HBSMCs were unstimulated (Unstim) or stimulated *in vitro* with 10 ng/mL recombinant human IL-1β in the presence of increasing concentrations of beta hydroxybutyric acid (BHBA), a combination of FFAR3 agonist (AR) and BHBA, or a combination of MCT1 inhibitor (AZD) and BHBA for 24 hours and **(A)** IL-6 or **(B)** CXCL8/IL-8 were measured. n=4 per group; values are inclusive of studies performed three times. ∗∗∗∗*P* ≤ 0.0001 compared to the vehicle, #*P* ≤ 0.05, ##*P* ≤ 0.01, ####*P* ≤ 0.0001 compared to IL-1β.

### FFAR3 Activation Is Sufficient to Inhibit IL-1β-Induced HBSMC Pro-Inflammatory Cytokine Production

Given that beta-hydroxybutyrate (BHB) has been suggested to act as a ligand for FFAR3 and exert its beneficial effects through a FFAR3-mediated pathway, we investigated whether activating FFAR3 could replicate the inhibitory effects of BHB on IL-1β-induced pro-inflammatory cytokine production from HBSMC. Whereas in our previous study we evaluated the combination of BHBA and FFAR3 agonist, we sought to determine whether FFAR3 activation alone could elicit the same effects. HBSMC were exposed to biologically relevant concentrations of BHBA, 50µM FFAR3 agonist (AR) alone, or to a combination of the concentrations of BHBA and 50µM FFAR3 agonist. HBSMCs exposed to BHBA and AR individually, as well as those exposed to the combination of BHBA and the FFAR3 agonist, significantly attenuated IL-1β-induced HBSMC production of IL-6 **(Figure 7A)** and CXCL8/IL-8 **(Figure 7B)**, suggesting that the observed inhibitory effects of BHB may indeed be mediated, at least in part, through the activation of FFAR3. Notably, whereas the FFAR3 agonist (AR) had a robust effect alone, the combination of the FFAR3 agonist and BHBA exerted the greatest inhibition, signifying a synergistic effect.

**Figure 7.**
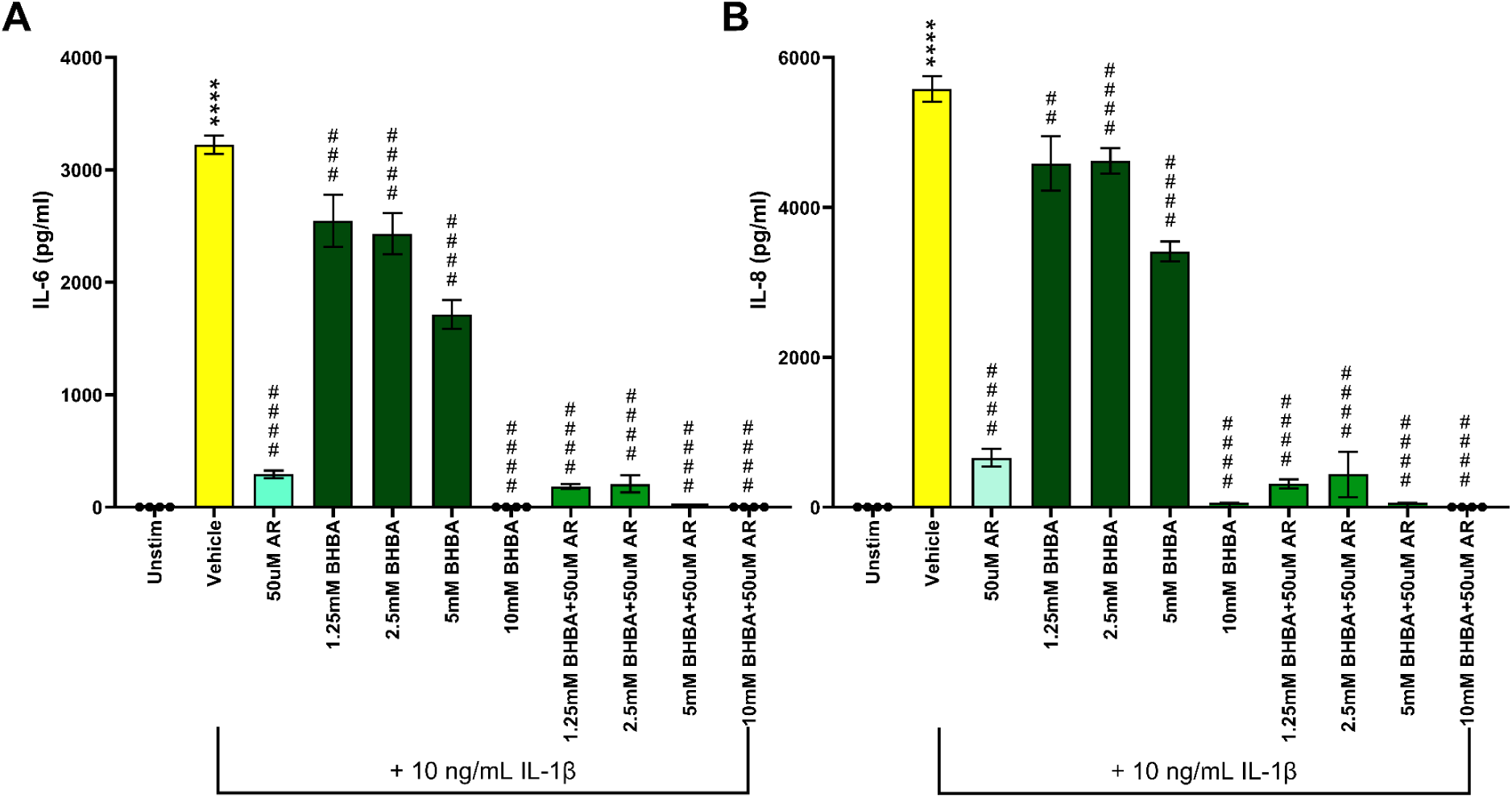
FFAR3 activation is sufficient to inhibit IL-1β induced pro-inflammatory cytokine secretion. HBSMCs were unstimulated (Unstim) or stimulated *in vitro* with 10 ng/mL recombinant human IL-1β in the presence of increasing concentrations of beta hydroxybutyric acid (BHBA), a combination of FFAR3 agonist (AR) and BHBA, or a combination of MCT1 inhibitor (AZD) and BHBA for 24 hours and **(A)** IL-6 or **(B)** CXCL8/IL-8 were measured. n=4 per group; values are inclusive of studies performed three times. ∗∗∗∗*P* ≤ 0.0001 compared to the vehicle, #*P* ≤ 0.05, ##*P* ≤ 0.01, ####*P* ≤ 0.0001 compared to IL-1β.

## Discussion

The growing asthma epidemic underscores the urgency for new treatment approaches aimed at managing severe and challenging endotypes, ultimately enhancing the well-being of individuals affected by this complex and chronic disorder. One promising strategy that has attracted increasing interest is therapeutic ketosis, recognized for its wide-ranging beneficial effects across various pathological conditions and diseases^53–56^. In our previous studies, we reported that therapeutic ketosis, achieved through dietary interventions such as a ketogenic diet, ketogenic precursor supplementation, or ketone ester intake, augments circulating concentrations of BHB and reduces methacholine hyperresponsiveness, a pathophysiological feature in models of preclinical allergic-associated and obesity-associated asthma^20,21^, decreases airway inflammation, and inhibits a multitude of pathological activities of asthma relevant cell types^20,21^. Several other studies have shown the *in vivo* anti-inflammatory activities of therapeutic ketosis^36,57,58^, with some affecting lung inflammation.

Although we previously demonstrated that exogenous ketones reduce pro-inflammatory cytokine production and exert anti-inflammatory effects *in vitro*^20,21^, their impact on bronchial smooth muscle cells remains unexplored. Given that bronchial smooth muscle cells are key players in bronchial inflammation, acting as both targets and mediators of inflammatory responses^5,6^, investigating their cytokine production represents a critical research area in airway inflammation and asthma management. In preclinical asthma models, distal airway and peripheral lung dysfunction are commonly observed^59^. Our prior work showed that elevated BHB concentrations confer protective effects on these regions ^20,21^, including reductions in methacholine-induced hyperresponsiveness, airway resistance, tissue damping, and tissue elastance—physiological markers that increase due to heterogeneous ventilation caused by uneven airway narrowing^59,60^ that are closely linked to peripheral airway contraction. These findings suggest that ketones may directly modulate bronchial smooth muscle to reduce airway hyperresponsiveness.

As reported herein, our studies demonstrate that BHB inhibits bronchial smooth muscle pro-inflammatory cytokine production induced by the asthma-relevant agonist, IL-1β^10,12,14–17^.

The mechanisms by which BHB can elicit these effects may include activation of cell surface receptors such as free fatty acid receptor 3 (FFAR3)^22,23,27,28^ or hydroxycarboxylic acid 2 (HCAR2)^25,26,29^. Given that BHB has been proposed to function across a wide range of applications and through many G-coupled protein receptors (GCPRs)^22,25–29^, including perhaps inducing tachyphylaxis, it likely operates through a fundamental mechanism that is universally relevant to these processes, such as ion exchange^27,36,61^, calcium signaling^27,28^, or altering membrane potential^61^.

Once determining that IL-1β significantly caused pro-inflammatory cytokines to be produced from human bronchial smooth muscle cells (HBSMC), our *in vitro* studies demonstrated that BHB attenuated the production of IL-6 and IL-8 (CXCL8) in a dose-dependent manner. These studies included conditions using beta-hydroxybutyric acid (BHBA) as well as sodium beta hydroxybutyrate (NaBHB). The inclusion of both of these compounds in our studies allowed us to explore the influence of pH by comparing BHBA with a 1M stock pH of 2.09 and the NaBHB, with a 1M stock pH of 9.73. When diluted to biologically relevant concentrations in buffered media, which resulted in near-neutral pH in the cell culture media (*i.e.*, pH 7.2), only BHBA had a significant inhibitory effect on IL-1β-induced production of pro-inflammatory cytokines *in vitro*. These results suggest that the acidic nature of the BHBA influences its ability to inhibit pro-inflammatory cytokine production. The inclusion of pH controls supports this notion in that when the 1M stock pH of the NaBHB was changed to be reminiscent of the 1M stock pH of the BHBA, the inhibitory effects of BHB were then present. Conversely, when the 1M stock pH of the BHBA was adjusted to the 1M stock pH of the NaBHB, the inhibitory effects were lost. The inclusion of a hydrochloric acid (HCl) pH-equivalent control, which did not inhibit pro-inflammatory cytokine production from IL-1β stimulated HBSMCs, implies that although the acid is sufficient to elicit modest inhibitory effects, the inclusion of BHB is necessary to enhance the effect.

BHBA is a racemic mixture containing both enantiomers (R)-BHB and (S)-BHB. Since only (R)-BHB can be efficiently metabolized by β-hydroxybutyrate dehydrogenase to form acetyl-CoA that is subsequently converted into ATP and used as an energy substrate^62^, we confirmed that the attenuation of pro-inflammatory cytokine production occurs independently of BHB’s metabolic function. Additionally, the MCT1 inhibitor, AZD3965 (AZD) failed to block BHB’s effects, indicating that MCT1-mediated uptake is not involved in the inhibitory capacity of BHB. The comparison between the (R)-BHB and the (R) enantiomer of the NaBHB (Na-R-BHB) reiterates that the acid allows for a robust pro-inflammatory cytokine production to be inhibited.

The HCAR2 agonists, nicotinic acid and sodium nicotinic acid, elicited similar inhibitory effects, but not to the extent of BHBA or the structurally similar compounds butyric acid and sodium butyrate, suggesting that some inhibitory effects may be mediated though HCAR2. Most likely, BHB does not elicit its attenuating effects on IL-1β-induced pro-inflammatory cytokine production through activation of HCAR2. Furthermore, the FFAR3 agonist AR420626 elicited inhibitory effects similar to and in an additive manner with BHB, suggesting that FFAR3 activation contributes substantially to the inhibitory action of BHB. Confirmation of this action would require an FFAR3 antagonist, a compound that is not currently available. Despite this limitation, these findings support future investigations, including studies using FFAR3 knockdown cells and FFAR3 knockout mice, to clarify the receptor’s involvement.

Simultaneous exposure to BHBA during IL-1β stimulation was more effective at reducing IL-1β-induced pro-inflammatory cytokine production in HBSMCs compared to BHBA pre-exposure and subsequent stimulation with IL-1β. This effect may be partly due to greater cytokine induction during simultaneous exposure, likely caused by increased cell confluency from the additional 24 hours in culture. Nonetheless, in a clinical setting, ‘therapeutic ketosis’ would maintain consistently elevated BHB concentrations, which could be modeled *in vitro* using the pre-exposure approach without removing BHBA before assessing IL-1β-induced pro-inflammatory cytokine production from HBSMCs.

There are several limitations to our findings. Namely, we conducted *in vitro* studies using primary human bronchial smooth muscle cells instead of *ex vivo* studies or human subjects. While informative, human cell studies fail to capture the prolonged and complex nature of human asthma. They offer a reductionist approach by focusing on a single cell type and a specific agonist that readily initiates an inflammatory cascade, allowing us to observe only the inhibitory effects of BHB at that initial step without assessing its role in maintaining suppression or facilitating recovery. Another notable limitation of this study is the lack of assessment of the fibrotic remodeling response of bronchial smooth muscle to IL-1β stimulation. In our previous work, we used an *in vivo* mouse model of severe asthma characterized by fibrotic remodeling and found that therapeutic ketosis still produced beneficial effects^21^. Overall, these findings support the potential of therapeutic ketosis as a promising treatment strategy for asthma, demonstrating its efficacy in both fibrotic and non-fibrotic contexts and highlighting its broad anti-inflammatory and protective effects. To gain a more comprehensive understanding of BHB’s impact on the long-term outcomes of an asthma exacerbation, diverse models, including *in vivo*, *ex vivo*, more advanced *in vitro* systems, and *in clinic* studies could be used to evaluate not only the initiation but also the maintenance and resolution of inflammation.

Therapeutic ketosis is being explored in the clinical treatment of respiratory diseases, including asthma^63,64^ and Cystic fibrosis^65^. In asthma trials, medium-chain triglyceride supplementation is used as a substrate to promote ketone body formation *in vivo*, whereas Cystic fibrosis trial uses a ketone ester precursor, which more rapidly and efficiently increases circulating ketone body concentrations and is the same compound we have employed in mouse models of obese asthma and allergic asthma^20,21^. The inclusion of therapeutic ketosis in clinical trials highlights the increasing scientific interest in its potential benefits. The positive results of the studies reported herein would strengthen the case for investigating therapeutic ketosis into asthma treatment regimens and stimulate further mechanistic and clinical studies into its capacity to serve as a complementary or alternative asthma therapy. This ongoing exploration emphasizes the importance of understanding ketone-based interventions and calls for continued examination of their underlying mechanisms and therapeutic potential.

## Graphical Abstract

In accordance with journal requirements, a graphical abstract will be submitted if we are invited to revise this manuscript.

## Abbreviations

AcAc: acetoacetate
ANOVA: analysis of variance
BHB: beta-hydroxybutyrate
FFAR: free fatty acid receptor
GPR: G protein-coupled receptor
HBSMC: human bronchial smooth muscle cells
HCAR: hydroxycarboxylic acid receptor
HDM: house dust mite
IL-: interleukin-
PBS: phosphate-buffered saline
MCT1: monocarboxylate transporter 1
SEM: standard error of the mean

## References

1. Statistics, N. C. f. H. Percentage of current asthma for adults aged 18 and over, United States, 2019—2023., (Center for Disease Control, 2023).

2 Cockcroft, D. W. Methacholine Challenge Testing in the Diagnosis of Asthma. Chest 158, 433–434 (2020). 10.1016/j.chest.2020.04.034

3 Wenzel, S. E. Asthma phenotypes: the evolution from clinical to molecular approaches. Nat Med 18, 716–725 (2012). 10.1038/nm.2678

4 Bara, I., Ozier, A., Tunon de Lara, J. M., Marthan, R. & Berger, P. Pathophysiology of bronchial smooth muscle remodelling in asthma. Eur Respir J 36, 1174–1184 (2010). 10.1183/09031936.00019810

5 Brightling, C. E. et al. Mast-cell infiltration of airway smooth muscle in asthma. N Engl J Med 346, 1699–1705 (2002). 10.1056/NEJMoa012705

6 Berger, P. et al. Tryptase-stimulated human airway smooth muscle cells induce cytokine synthesis and mast cell chemotaxis. FASEB J 17, 2139–2141 (2003). 10.1096/fj.03-0041fje

7 Fahy, J. V. Type 2 inflammation in asthma--present in most, absent in many. Nat Rev Immunol 15, 57–65 (2015). 10.1038/nri3786

8 Peebles, R. S., Jr. Is IL-1beta inhibition the next therapeutic target in asthma? J Allergy Clin Immunol 139, 1788–1789 (2017). 10.1016/j.jaci.2017.03.018

9 McKay, S. & Sharma, H. S. Autocrine regulation of asthmatic airway inflammation: role of airway smooth muscle. Respir Res 3, 11 (2002). 10.1186/rr160

10 Shea-Donohue, T., Notari, L., Sun, R. & Zhao, A. Mechanisms of smooth muscle responses to inflammation. Neurogastroenterol Motil 24, 802–811 (2012). 10.1111/j.1365-2982.2012.01986.x

11 Yamauchi, K. & Ogasawara, M. The Role of Histamine in the Pathophysiology of Asthma and the Clinical Efficacy of Antihistamines in Asthma Therapy. Int J Mol Sci 20 (2019). 10.3390/ijms20071733

12 Martin, J. G. [Animal models of bronchial hyperreactivity]. Rev Mal Respir 11, 93–99 (1994).

13 Elias, J. A., Zhu, Z., Chupp, G. & Homer, R. J. Airway remodeling in asthma. J Clin Invest 104, 1001–1006 (1999). 10.1172/JCI8124

14 Pena-Garcia, P. E. et al. Bariatric surgery decreases the capacity of plasma from obese asthmatic subjects to augment airway epithelial cell proinflammatory cytokine production. Am J Physiol Lung Cell Mol Physiol 326, L71–L82 (2024). 10.1152/ajplung.00205.2023

15 Liao, Z. et al. IL-1beta: a key modulator in asthmatic airway smooth muscle hyper-reactivity. Expert Rev Respir Med 9, 429–436 (2015). 10.1586/17476348.2015.1063422

16 Fu, J. J., McDonald, V. M., Baines, K. J. & Gibson, P. G. Airway IL-1beta and Systemic Inflammation as Predictors of Future Exacerbation Risk in Asthma and COPD. Chest 148, 618–629 (2015). 10.1378/chest.14-2337

17 Thomas, S. S. & Chhabra, S. K. A study on the serum levels of interleukin-1beta in bronchial asthma. J Indian Med Assoc 101, 282, 284, 286 passim (2003).

18 Gans, M. D. & Gavrilova, T. Understanding the immunology of asthma: Pathophysiology, biomarkers, and treatments for asthma endotypes. Paediatr Respir Rev 36, 118–127 (2020). 10.1016/j.prrv.2019.08.002

19 Mims, J. W. Asthma: definitions and pathophysiology. Int Forum Allergy Rhinol 5 Suppl 1, S2–6 (2015). 10.1002/alr.21609

20 Mank, M. M. et al. Therapeutic ketosis decreases methacholine hyperresponsiveness in mouse models of inherent obese asthma. Am J Physiol Lung Cell Mol Physiol 322, L243–L257 (2022). 10.1152/ajplung.00309.2021

21 Mank, M. M. et al. Ketone body augmentation decreases methacholine hyperresponsiveness in mouse models of allergic asthma. J Allergy Clin Immunol Glob 1, 282–298 (2022). 10.1016/j.jacig.2022.08.001

22 Puchalska, P. & Crawford, P. A. Multi-dimensional Roles of Ketone Bodies in Fuel Metabolism, Signaling, and Therapeutics. Cell Metab 25, 262–284 (2017). 10.1016/j.cmet.2016.12.022

23 Puchalska, P. & Crawford, P. A. Metabolic and Signaling Roles of Ketone Bodies in Health and Disease. Annu Rev Nutr 41, 49–77 (2021). 10.1146/annurev-nutr-111120-111518

24 Murakami, M. & Tognini, P. Molecular Mechanisms Underlying the Bioactive Properties of a Ketogenic Diet. Nutrients 14 (2022). 10.3390/nu14040782

25 Chen, Y. et al. beta-Hydroxybutyrate protects from alcohol-induced liver injury via a Hcar2-cAMP dependent pathway. J Hepatol 69, 687–696 (2018). 10.1016/j.jhep.2018.04.004

26 Qi, J. et al. Beta-Hydroxybutyrate: A Dual Function Molecular and Immunological Barrier Function Regulator. Front Immunol 13, 805881 (2022). 10.3389/fimmu.2022.805881

27 Newman, J. C. & Verdin, E. beta-Hydroxybutyrate: A Signaling Metabolite. Annu Rev Nutr 37, 51–76 (2017). 10.1146/annurev-nutr-071816-064916

28 Won, Y. J., Lu, V. B., Puhl, H. L., 3rd & Ikeda, S. R. beta-Hydroxybutyrate modulates N-type calcium channels in rat sympathetic neurons by acting as an agonist for the G-protein-coupled receptor FFA3. J Neurosci 33, 19314–19325 (2013). 10.1523/JNEUROSCI.3102-13.2013

29 Thio, C. L., Lai, A. C., Ting, Y. T., Chi, P. Y. & Chang, Y. J. The ketone body beta-hydroxybutyrate mitigates ILC2-driven airway inflammation by regulating mast cell function. Cell Rep 40, 111437 (2022). 10.1016/j.celrep.2022.111437

30 Li, R. et al. Adaptive Metabolic Responses Facilitate Blood-Brain Barrier Repair in Ischemic Stroke via BHB-Mediated Epigenetic Modification of ZO-1 Expression. Adv Sci (Weinh) 11, e2400426 (2024). 10.1002/advs.202400426

31 Lu, Y. et al. Ketogenic diet attenuates oxidative stress and inflammation after spinal cord injury by activating Nrf2 and suppressing the NF-kappaB signaling pathways. Neurosci Lett 683, 13–18 (2018). 10.1016/j.neulet.2018.06.016

32 Haces, M. L. et al. Antioxidant capacity contributes to protection of ketone bodies against oxidative damage induced during hypoglycemic conditions. Exp Neurol 211, 85–96 (2008). 10.1016/j.expneurol.2007.12.029

33 Shimazu, T. et al. Suppression of oxidative stress by beta-hydroxybutyrate, an endogenous histone deacetylase inhibitor. Science 339, 211–214 (2013). 10.1126/science.1227166

34 Hoyt, L. R. et al. Ethanol and Other Short-Chain Alcohols Inhibit NLRP3 Inflammasome Activation through Protein Tyrosine Phosphatase Stimulation. J Immunol 197, 1322–1334 (2016). 10.4049/jimmunol.1600406

35 Theofani, E., Semitekolou, M., Morianos, I., Samitas, K. & Xanthou, G. Targeting NLRP3 Inflammasome Activation in Severe Asthma. J Clin Med 8 (2019). 10.3390/jcm8101615

36 Youm, Y. H. et al. The ketone metabolite beta-hydroxybutyrate blocks NLRP3 inflammasome-mediated inflammatory disease. Nat Med 21, 263–269 (2015). 10.1038/nm.3804

37 Johnson, J. B. et al. Alternate day calorie restriction improves clinical findings and reduces markers of oxidative stress and inflammation in overweight adults with moderate asthma. Free Radic Biol Med 42, 665–674 (2007). 10.1016/j.freeradbiomed.2006.12.005

38 Jensen, M. E., Gibson, P. G., Collins, C. E., Hilton, J. M. & Wood, L. G. Diet-induced weight loss in obese children with asthma: a randomized controlled trial. Clin Exp Allergy 43, 775–784 (2013). 10.1111/cea.12115

39 Umemura, A. et al. Relationships Between Changes in Serum Ketone Body Levels and Metabolic Effects in Patients with Severe Obesity Who Underwent Laparoscopic Sleeve Gastrectomy. Obesity surgery 34, 2607–2616 (2024). 10.1007/s11695-024-07337-8

40 Aberle, J. et al. Metformin after bariatric surgery--an acid problem. Exp Clin Endocrinol Diabetes 120, 152–153 (2012). 10.1055/s-0031-1285911

41 Angelidi, A. M. et al. Early metabolomic, lipid and lipoprotein changes in response to medical and surgical therapeutic approaches to obesity. Metabolism: clinical and experimental 138, 155346 (2023). 10.1016/j.metabol.2022.155346

42 Stefan, M., Sharp, M., Gheith, R., Lowery, R. & Wilson, J. The Effect of Exogenous Beta-Hydroxybutyrate Salt Supplementation on Metrics of Safety and Health in Adolescents. Nutrients 13 (2021). 10.3390/nu13030854

43 Hashim, S. A. & VanItallie, T. B. Ketone body therapy: from the ketogenic diet to the oral administration of ketone ester. J Lipid Res 55, 1818–1826 (2014). 10.1194/jlr.R046599

44 Gosens, R., Zaagsma, J., Meurs, H. & Halayko, A. J. Muscarinic receptor signaling in the pathophysiology of asthma and COPD. Respir Res 7, 73 (2006). 10.1186/1465-9921-7-73

45 Fryer, A. D. & Jacoby, D. B. Muscarinic receptors and control of airway smooth muscle. American journal of respiratory and critical care medicine 158, S154–160 (1998). 10.1164/ajrccm.158.supplement_2.13tac120

46 Mak, J. C. & Barnes, P. J. Autoradiographic visualization of muscarinic receptor subtypes in human and guinea pig lung. Am Rev Respir Dis 141, 1559–1568 (1990). 10.1164/ajrccm/141.6.1559

47 Eglen, R. M., Hegde, S. S. & Watson, N. Muscarinic receptor subtypes and smooth muscle function. Pharmacol Rev 48, 531–565 (1996).

48 Schmidt, D. & Rabe, K. F. Immune mechanisms of smooth muscle hyperreactivity in asthma. J Allergy Clin Immunol 105, 673–682 (2000). 10.1067/mai.2000.105705

49 Shivva, V. et al. The Population Pharmacokinetics of D-beta-hydroxybutyrate Following Administration of (R)-3-Hydroxybutyl (R)-3-Hydroxybutyrate. AAPS J 18, 678–688 (2016). 10.1208/s12248-016-9879-0

50 Vinolo, M. A., Rodrigues, H. G., Nachbar, R. T. & Curi, R. Regulation of inflammation by short chain fatty acids. Nutrients 3, 858–876 (2011). 10.3390/nu3100858

51 Du, Y. et al. The Role of Short Chain Fatty Acids in Inflammation and Body Health. Int J Mol Sci 25 (2024). 10.3390/ijms25137379

52 Qiu, Y. et al. Ketogenic diet alleviates renal fibrosis in mice by enhancing fatty acid oxidation through the free fatty acid receptor 3 pathway. Front Nutr 10, 1127845 (2023). 10.3389/fnut.2023.1127845

53 Daines, S. A. The Therapeutic Potential and Limitations of Ketones in Traumatic Brain Injury. Front Neurol 12, 723148 (2021). 10.3389/fneur.2021.723148

54 Waldman, H. S. & McAllister, M. J. Exogenous Ketones as Therapeutic Signaling Molecules in High-Stress Occupations: Implications for Mitigating Oxidative Stress and Mitochondrial Dysfunction in Future Research. Nutr Metab Insights 13, 1178638820979029 (2020). 10.1177/1178638820979029

55 Yurista, S. R. et al. Therapeutic Potential of Ketone Bodies for Patients With Cardiovascular Disease: JACC State-of-the-Art Review. J Am Coll Cardiol 77, 1660–1669 (2021). 10.1016/j.jacc.2020.12.065

56 Zarnowska, I. M. Therapeutic Use of the Ketogenic Diet in Refractory Epilepsy: What We Know and What Still Needs to Be Learned. Nutrients 12 (2020). 10.3390/nu12092616

57 Goldberg, E. L. et al. Ketogenic diet activates protective gammadelta T cell responses against influenza virus infection. Sci Immunol 4 (2019). 10.1126/sciimmunol.aav2026

58 Ryu, S. et al. Ketogenic diet restrains aging-induced exacerbation of coronavirus infection in mice. Elife 10 (2021). 10.7554/eLife.66522

59 Bates, J. Lung Mechanics: An Inverse Modeling Approach.. 169–187 (2009).

60 Wagers, S., Lundblad, L. K., Ekman, M., Irvin, C. G. & Bates, J. H. The allergic mouse model of asthma: normal smooth muscle in an abnormal lung? Journal of applied physiology 96, 2019–2027 (2004). 10.1152/japplphysiol.00924.2003

61 Manville, R. W., Papanikolaou, M. & Abbott, G. W. M-Channel Activation Contributes to the Anticonvulsant Action of the Ketone Body beta-Hydroxybutyrate. J Pharmacol Exp Ther 372, 148–156 (2020). 10.1124/jpet.119.263350

62 Webber, R. J. & Edmond, J. Utilization of L(+)-3-hydroxybutyrate, D(-)-3-hydroxybutyrate, acetoacetate, and glucose for respiration and lipid synthesis in the 18-day-old rat. J Biol Chem 252, 5222–5226 (1977).

63 Georas, S. N. et al. The Precision Interventions for Severe and/or Exacerbation-Prone (PrecISE) Asthma Network: An overview of Network organization, procedures, and interventions. J Allergy Clin Immunol 149, 488–516 e489 (2022). 10.1016/j.jaci.2021.10.035

64 ClincialTrials.gov. Identifier: NCT06151405, Nutrition for Asthmatics (INHALE). National Library of Medicine (US) (2024).

65 Plaisance, E. P. et al. Low-Dose Ketone Monoester Administration in Adults with Cystic Fibrosis: A Pilot and Feasibility Study. Nutrients 16 (2024). 10.3390/nu16223957

